# Parallel processing of orthogonal manifolds enables zero-shot composition in recurrent networks

**DOI:** 10.64898/2026.06.14.732142

**Authors:** Yuma Osako, Aineias Arango, Toshitake Asabuki

## Abstract

Animals flexibly combine learned behaviors into novel actions without practicing their combinations, yet the computational mechanisms that enable independently acquired computations to be expressed in parallel remain unclear. Here we show that feedback geometry during learning determines whether recurrent dynamics can be recombined through zero-shot parallel composition. Using recurrent networks trained by a local predictive plasticity rule, we found that distinct feedback vectors embed independently learned computations in separable dynamical subspaces, allowing novel input combinations to co-activate these components and generate composite outputs without joint training. In contrast, aligned feedback vectors, as well as networks trained by backpropagation through time, exhibited accurate single-task performance but failed to support parallel composition, demonstrating that task acquisition and future reusability are dissociable properties of learning. A combined input evoked a single composite population trajectory, whose projections onto feedback-shaped task subspaces recovered the independently learned component dynamics. The same principle reproduced additive reach-posture geometry observed in motor cortex and generalized to higher-dimensional movement primitives. These results identify feedback geometry as a computational principle by which learning systems structure recurrent dynamics for future compositional reuse.

## Introduction

Animals exhibit remarkable behavioral flexibility, enabling them to rapidly adapt to novel situations without learning every possible behavioral contingency^1^. Such flexibility requires more than accurate acquisition of individual behavioral components. It requires that learned computations remain available for simultaneous expression when new combinations are encountered^2,3^. A neural circuit may acquire each component accurately yet embed them in a geometry that prevents their interference^4–7^. Task acquisition and future compositionality are therefore distinct computational properties.

Population geometry offers a natural framework for understanding this distinction^8^. Individual neurons often exhibit multiplexed responses to different task variables^9,10^, yet population activity often represents these variables in separable or nearly orthogonal subspaces^5–7,11–14^, allowing multiple signals to coexist within a shared population. In motor cortex, for example, movement-related dynamics and posture-related activity occupy separable subspaces, so that changes in posture shift the reach manifold without disrupting its internal dynamical structure^15^. Similar forms of separable geometry have been observed for task variables^6,12,16^, contexts^17,18^, preparatory states^19^, and movement-related dynamics^20,21^ across cortical areas. These findings suggest that neural manifolds may provide a substrate for flexible computation. However, most studies have characterized population geometry after learning is complete. How plasticity creates geometries that support not only individual task performance, but also future composition, remains unclear.

This problem is difficult to isolate in conventional models of compositionality. Many models achieve combination through modular architectures, explicit factorization, joint multitask training, baseline control, or direct exposure to combined conditions^2,3,22,23^. In these settings, the geometry needed for combination is either imposed by design or shaped by experience with the relevant combinations. Biological circuits, however, often acquire behavioral components across separate learning episodes and later encounter novel situations in which those components must be expressed together^24,25^. Moreover, compositionality is often studied in settings where components are arranged sequentially over time^2,3,26,27^, whereas many behaviors require multiple computations to be expressed in parallel^28^, as when movement goals and body posture must be represented simultaneously during reaching. Whether a recurrent circuit can support such zero-shot composition therefore depends on how independently acquired computations are embedded during learning.

Here we study this problem as zero-shot parallel composition, in which multiple learned computations must be expressed simultaneously within the same recurrent population. This setting isolates a general requirement for compositional learning: acquired dynamics must not only solve the tasks on which they were trained, but also remain structured so that they can be reused in novel combinations. Using recurrent networks trained by a local predictive plasticity rule, we show that task-specific feedback vectors embed independently acquired dynamics in separable subspaces that preserve them for later simultaneous expression. Combined inputs can then co-activate these feedback-shaped dynamical components and produce composite outputs without training on the combined condition. By contrast, aligned feedback vectors or standard backpropagation through time allow individual task acquisition but fail to support zero-shot composition. We establish this principle in an abstract temporal-generation task, identify a population-level mechanism based on projections onto feedback-shaped task subspaces, and show that the same mechanism reproduces motor cortex-like reach-posture geometry and extends to higher-dimensional multi-output movement-like primitives.

## Results

### Zero-shot parallel composition of independently learned primitives

We first define the computational problem of zero-shot parallel composition. As an intuitive example, a dancer may learn one movement component, such as spinning, and separately learn another, such as jumping (Fig. 1A). Zero-shot parallel composition asks whether these learned components can later be expressed together, as in a spinning jump, without prior experience of that combination (Fig. 1B).

**Figure 1.**
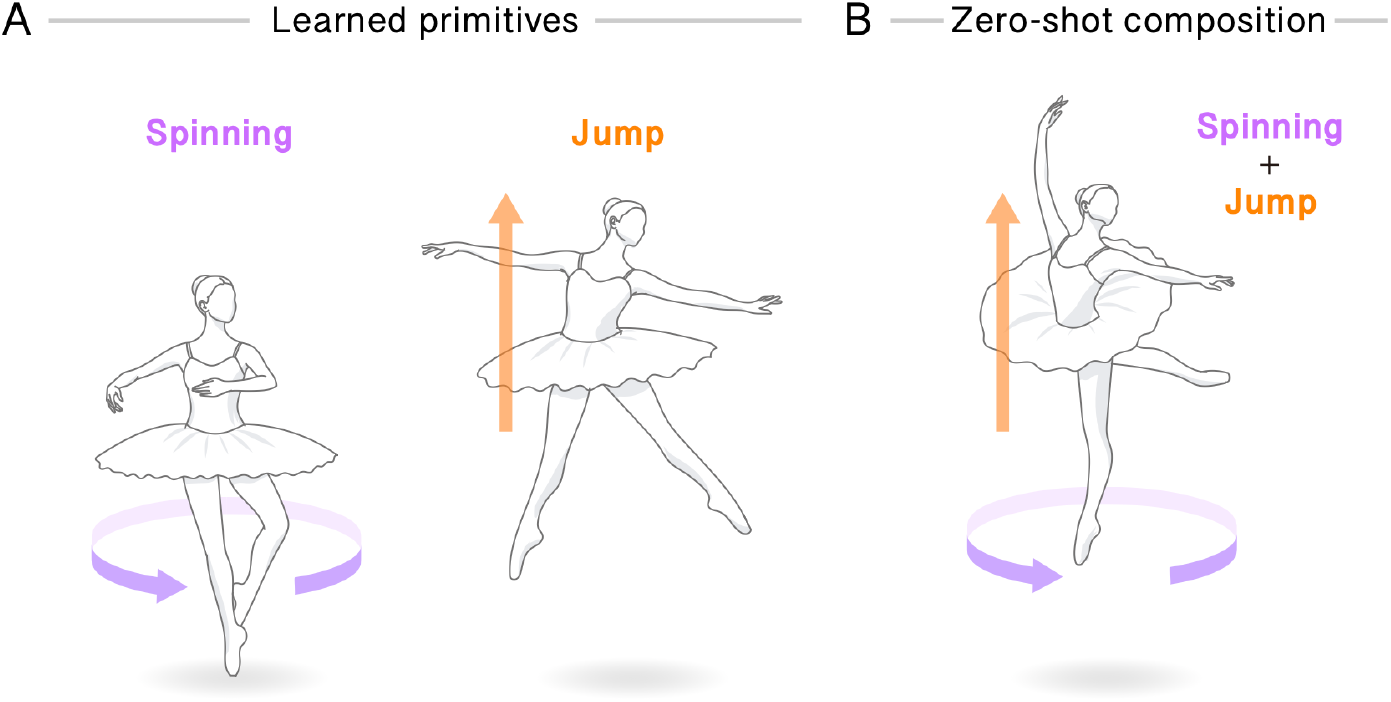
Zero-shot parallel composition without joint training. **A** A dancer separately learns a spinning movement and jump. **B** Zero-shot parallel composition asks whether the two independently acquired components can later be expressed together without practicing the combined behavior.

More generally, we considered a recurrent network that learns two distinct computations in separate training blocks and is then tested on a novel condition requiring both computations to be performed simultaneously. Because the network is never exposed to this combined condition during training, successful performance requires the learned computations to remain reusable beyond their original training conditions. This setting differs from conventional multitask learning, in which networks may be trained directly on task combinations, and from sequential composition, where computations unfold over time rather than being expressed in parallel.

### Feedback geometry determines zero-shot composition

We reasoned that a recurrent circuit could support such composition if independently learned computations were embedded into orthogonal subspaces of population activity. Such an organization would preserve each computation while allowing multiple learned dynamics to be expressed in parallel within the same recurrent population. Building on previous work showing that distinct feedback signals can organize learned dynamics into orthogonal task subspaces^29^, we tested whether feedback geometry determines whether independently learned computations remain reusable during zero-shot composition.

To this end, we used a sine-wave generation task in which a recurrent network received one of two distinct constant inputs, S1 or S2 (Fig. 2A). Each input was associated with a different sinusoidal target output. The two input-output mappings were trained separately, and the network was never exposed to trials in which both inputs were presented simultaneously. Learning was implemented using a local predictive plasticity rule in which feedback signals constrained recurrent synaptic updates through a locally computed prediction error^30^. We compared two conditions: a different-feedback condition, in which each task was associated with a different feedback vector (Diff FB), and a same-feedback (Same FB) condition, in which both tasks shared an identical feedback vector. Although learning was somewhat slower in the Same FB condition, both conditions eventually acquired the two individual tasks with high accuracy (Supplementary Fig. 1). This allowed us to ask whether feedback geometry affects zero-shot composition beyond its effect on acquiring the individual tasks.

**Figure 2.**
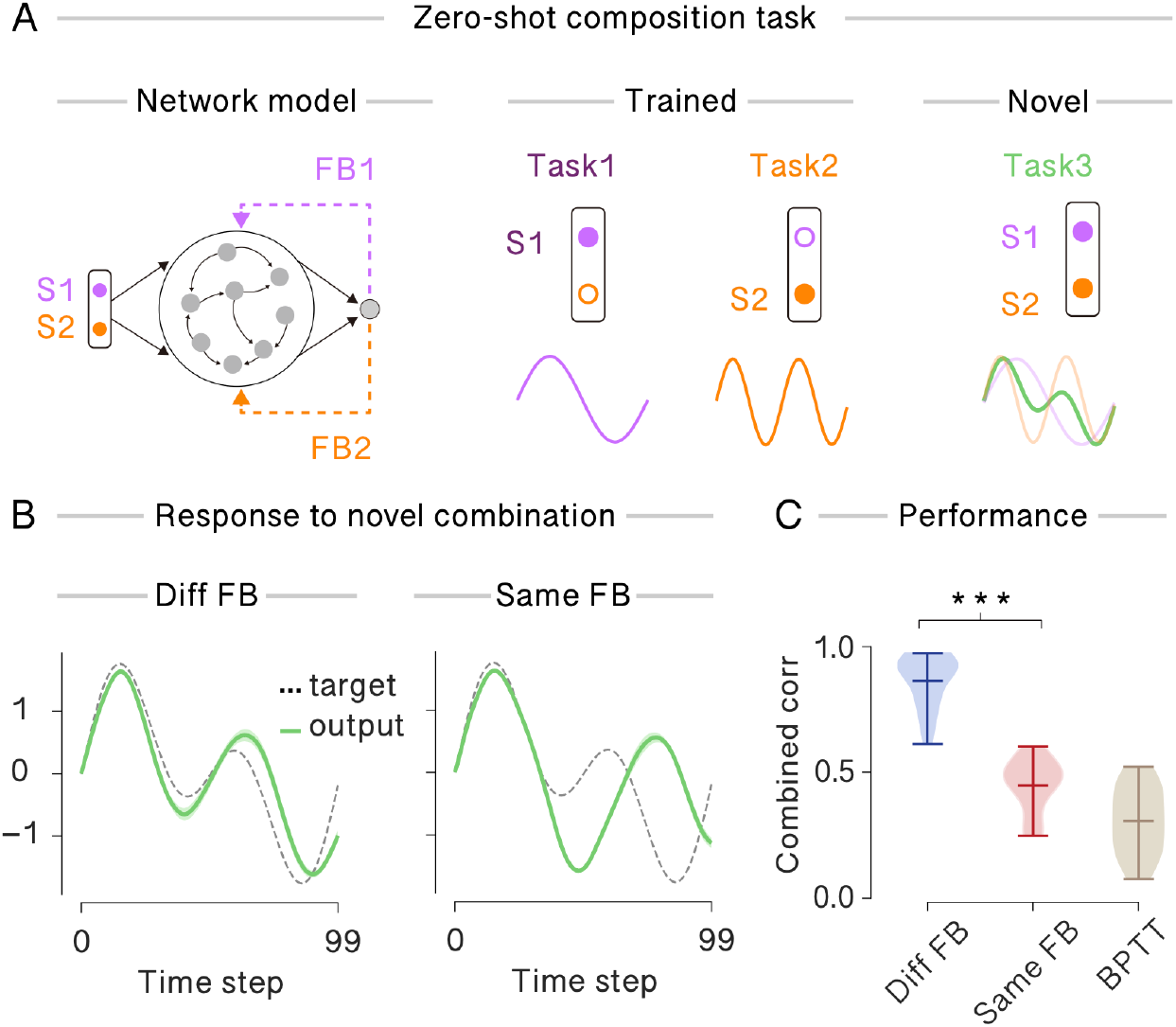
Feedback geometry determines zero-shot composition. **A** Minimal temporal-generation task. A recurrent network receives two input signals, S1 and S2, and generates a single output. The two tasks were trained separately, with each input associated with a different sinusoidal target output. After training, both inputs were presented simultaneously, and the network output was compared with an untrained combined target defined as the sum of the two individual targets. **B** Combined-condition output. In the different-feedback condition, in which the two tasks were trained with distinct feedback vectors, the network output closely followed the combined target. In the same-feedback condition, in which both tasks were trained with the same feedback vector, the network failed to reproduce the combined target. **C** Output correlation with the combined target across networks. Networks trained with different feedback vectors showed significantly higher combined-condition performance than networks trained with the same feedback vector. BPTT-trained networks showed comparable combined-condition performance to the same-feedback condition. ***p < 0.001, Diff FB vs Same FB.

After training, we evaluated zero-shot composition by presenting S1 and S2 simultaneously. The target output was defined as the linear sum of the two individually learned outputs. Successful performance therefore required the network to represent both learned computations concurrently through the same recurrent circuitry and readout. Networks trained with different feedback vectors closely tracked the combined target when S1 and S2 were presented simultaneously, indicating successful zero-shot composition (Fig. 2B; Diff FB). In contrast, networks trained with an identical feedback vector successfully learned the individual tasks but failed to reproduce the combined target under simultaneous input (Fig. 2B; Same FB). Across networks, correlation between output and the combined target was significantly higher when feedback vectors were distinct than when they were fully aligned (Fig. 2C).

We also confirmed recurrent networks trained with backpropagation through time (BPTT) learned the individual mappings but generalized poorly to the combined condition (Supplementary Fig. 2), performing similarly to networks trained with same-feedback condition (Fig. 2C). Thus, successful zero-shot composition did not follow automatically from single-task acquisition. Instead, it depended on how learning embedded the component computations in recurrent state space.

Together, these results indicate that feedback geometry is crucial to determine whether independently acquired computations remain available for later composition. Distinct feedback vectors enabled the network to reuse learned computations under a novel combined input, whereas aligned feedback supported individual task learning but impaired their simultaneous expression.

### A single composite trajectory recovers independently learned dynamics

We next asked how feedback geometry enables a single recurrent network to express two independently learned computations simultaneously. If distinct feedback signals embed the corresponding computational dynamics into different subspaces of the network, each computation may remain independently executable even when the inputs are combined. To test this possibility, we defined each task-specific subspace as the leading 10 principal components of the recurrent activity evoked by the corresponding single-input condition (Methods). We then examined how network activity under the combined-input condition was represented within these task-specific subspaces.

Under the different-feedback condition, the combined input drove the corresponding task-specific dynamics within each subspace. Consistent with this separability, the task-specific subspaces exhibited high principal angles (Supplementary Fig. 3). In the S1 subspace, network activity closely followed the trajectory evoked by S1 alone. Similarly, in the S2 subspace, activity recapitulated the dynamics evoked by S2 alone (Fig. 3A). Thus, simultaneous presentation of S1 and S2 activated the two learned dynamical processes in parallel within distinct subspaces. The resulting output was generated through the combination of these concurrently expressed computations.

**Figure 3.**
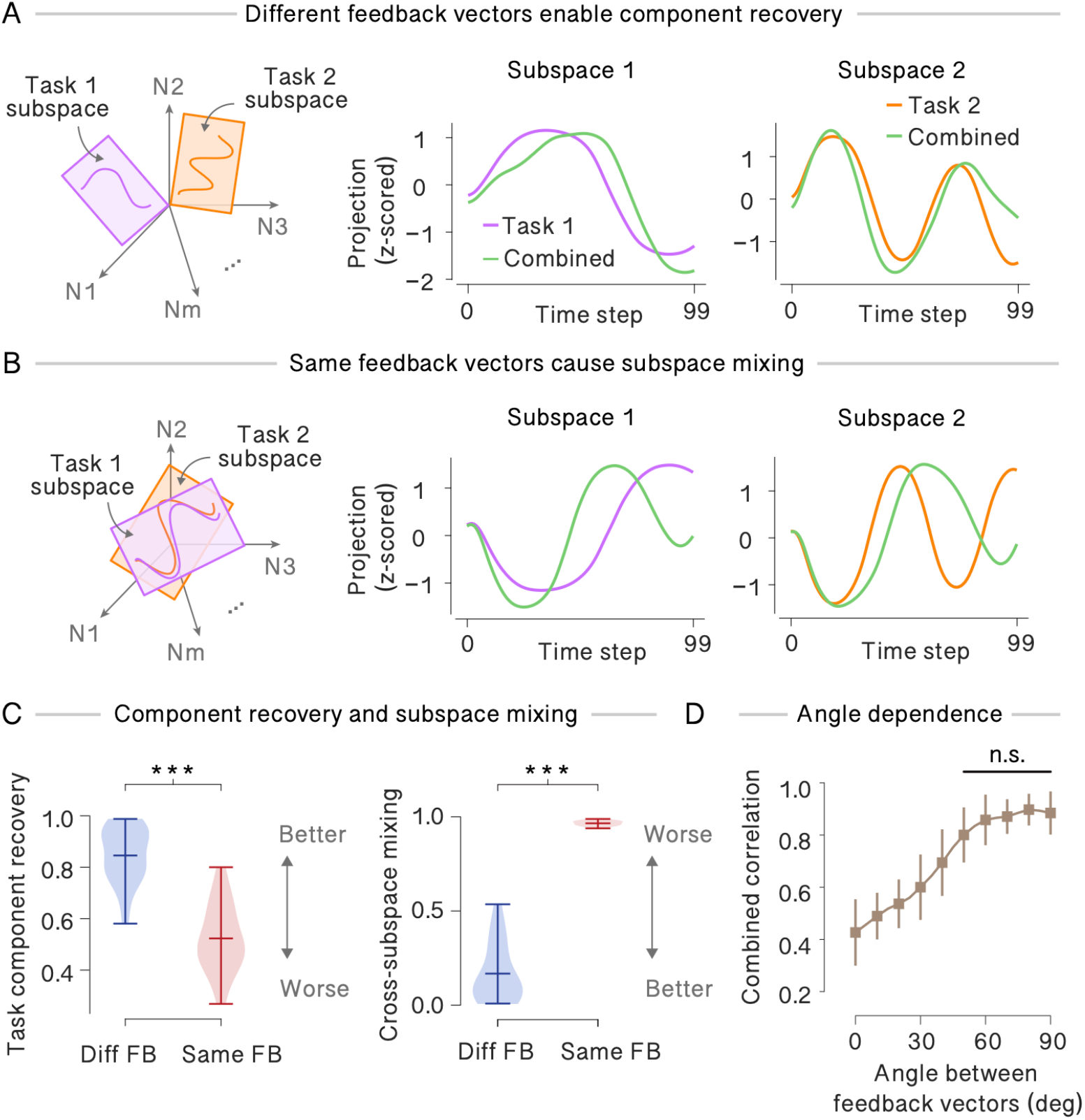
A single composite trajectory recovers independently learned component dynamics. **A** Distinct feedback enables component recovery. The combined-input trajectory was projected onto the subspace defined by each single-task trajectory. In the S1 subspace, the combined projection closely followed the S1-alone trajectory. In the S2 subspace, it closely followed the S2-alone trajectory, indicating that both learned component dynamics were recovered from a single composite population trajectory. **B** Shared feedback causes subspace mixing. The combined trajectory failed to preserve task-specific structure in either subspace, indicating that the learned components could not be simultaneously recovered. **C** Component recovery and subspace mixing. Left, component recovery quantified as the correlation between each single-task trajectory and the corresponding projection of the combined trajectory. Right, subspace mixing quantified as the absolute correlation between the combined-input projections onto the two task subspaces. **D** Feedback-angle dependence. The angle between the two feedback vectors was systematically varied. Zero-shot composition performance remained high when feedback vectors were sufficiently separated and declined as they became aligned. Comparisons against the 90° condition were not significant for sufficiently separated feedback vectors (n.s., p > 0.05, two-sample t-tests with Holm-Bonferroni correction for multiple comparisons).

In contrast, under the same-feedback condition, the two tasks occupied highly overlapping subspaces, as reflected by lower principal angles between the corresponding task subspaces. (Supplementary Fig. 3). Although the network successfully learned each task individually, the combined input drove activity within a shared subspace, causing the resulting trajectories to deviate substantially from those observed during the corresponding single-task conditions (Fig. 3B). Thus, the failure of zero-shot composition did not arise from an inability to acquire the component computations themselves. Rather, it resulted from a representational geometry in which distinct computations were not embedded into separate subspaces, leading to interference when both computations were simultaneously engaged.

We quantified the extent to which each task-specific dynamical component was preserved within its corresponding subspace under the combined-input condition. For each task subspace, we computed the correlation between the population trajectory evoked by the combined input and the corresponding trajectory observed during the single-task condition. These correlations were significantly higher in the different-feedback condition for both task subspaces (Fig. 3C; left), indicating that the original task-specific dynamics were more preserved during composition. In addition, cross-correlation between the activity projected onto the two task subspaces was significantly lower in the different-feedback condition (Fig. 3C; right), suggesting reduced crosstalk between the co-expressed dynamical processes.

We next asked whether the geometric relationship between feedback signals continuously controls compositional performance. To test this, we systematically varied the angle between the two feedback vectors (Fig. 3D). As the feedback vectors became more orthogonal, the network exhibited progressively better zero-shot composition. Performance remained high across a broad range of angles (approximately 60-90°) but declined as the two vectors became increasingly aligned. Thus, exact orthogonality was not required, yet the feedback vectors needed to be sufficiently distinct to prevent interference between learned subspaces.

In summary, population-level decomposition of the combined responses revealed the mechanism underlying zero-shot composition. Successful performance on the novel combined-input condition did not require additional training. Instead, each learned computation was preserved within a feedback-defined task subspace, and the combined response emerged through the concurrent expression and integration of these task-specific dynamical components. These results show that feedback geometry shapes recurrent activity into separable subspaces that can be co-activated during composition.

### Low-rank collective dynamics explain feedback-dependent composition

We next asked how orthogonal subspaces enables parallel composition. To gain mechanistic insight, we developed a low-rank mean-field description of the trained recurrent network^31^. Previous work has shown that feedback-guided learning can embed task-specific dynamics into structured recurrent modes^21^. Motivated by this, we approximated the trained recurrent connectivity as the sum of a random recurrent component and two task-specific low-rank modes,

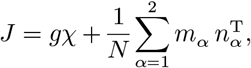

where *gχ* denotes the random recurrent component, and *m*_*α*_ and *n*_*α*_ are the right and left connectivity vectors associated with task α. In this framework, each learned mode can then be summarized by a low-dimensional collective variable,

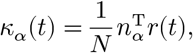

which quantifies how strongly the population activity engages the corresponding learned mode.

Linearizing the dynamics of these collective variables around the trained trajectory gives an effective two-dimensional dynamical system,

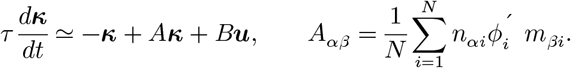

The full derivation of this reduced description, including the mean-field approximation and linearization, is provided in the Supplementary Information. In this framework, the diagonal terms *A*_11_ and *A*_22_ describe the intrinsic dynamics of the two learned computations, whereas the off-diagonal terms *A*_12_ and *A*_21_ quantify cross-coupling between them. Thus, successful zero-shot composition requires the effective dynamics to be approximately decoupled across modes.

When feedback vectors are distinct, learning embeds the two tasks into statistically separated recurrent modes. Consequently, the cross-coupling terms vanish up to finite-size fluctuations,

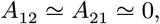

yielding an approximately decoupled dynamical system. The two collective variables can therefore be co-activated independently under combined input, and if the readout is aligned with the learned modes, the network response becomes approximately additive,

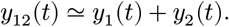

By contrast, aligned feedback vectors produce overlapping learned modes and finite cross-coupling. Activation of one computation then perturbs the dynamics of the other, preventing a simple superposition of the two single-task responses despite accurate acquisition of each task individually. This mean-field theory provides a mechanistic explanation for both the zero-shot composition under distinct feedback and its failure under shared feedback. It also explains the graded dependence on feedback angle observed in Fig. 3D. As the feedback angle between feedback vectors decreases, the learned modes overlap more strongly, strengthening cross-coupling and consequently impairing composition.

### Feedback-guided learning reproduces additive reach-posture geometry observed in motor cortex

Having established the mechanism for parallel processing in an abstract temporal-generation task, we next asked whether feedback-driven learning could account for geometric structures previously observed in motor cortex. During reaching from different initial postures, movement-related dynamics and posture-related activity occupy approximately separable subspaces, such that posture shifts the reach-related trajectories without distorting their internal dynamical structure^15^. We therefore asked whether a similar additive representational structure could emerge through separate learning of movement and posture, without exposure to their combinations.

We trained recurrent networks to learn movement and posture readouts in separate tasks, with each task associated with its own output and feedback signal (Fig. 4A, B). In the movement task, the network generated two-dimensional reach trajectories toward one of eight target directions. In the posture task, the network maintained one of three scalar posture outputs. During training, movement and posture inputs were never presented simultaneously, and distinct feedback vectors guided learning for the movement and posture readouts. The trained network successfully acquired both component tasks (Fig. 4C). During movement-only trials, the movement readouts generated direction-specific reach trajectories, whereas during posture-only trials, the posture readout converged to the corresponding target level while movement outputs remained near zero.

**Figure 4.**
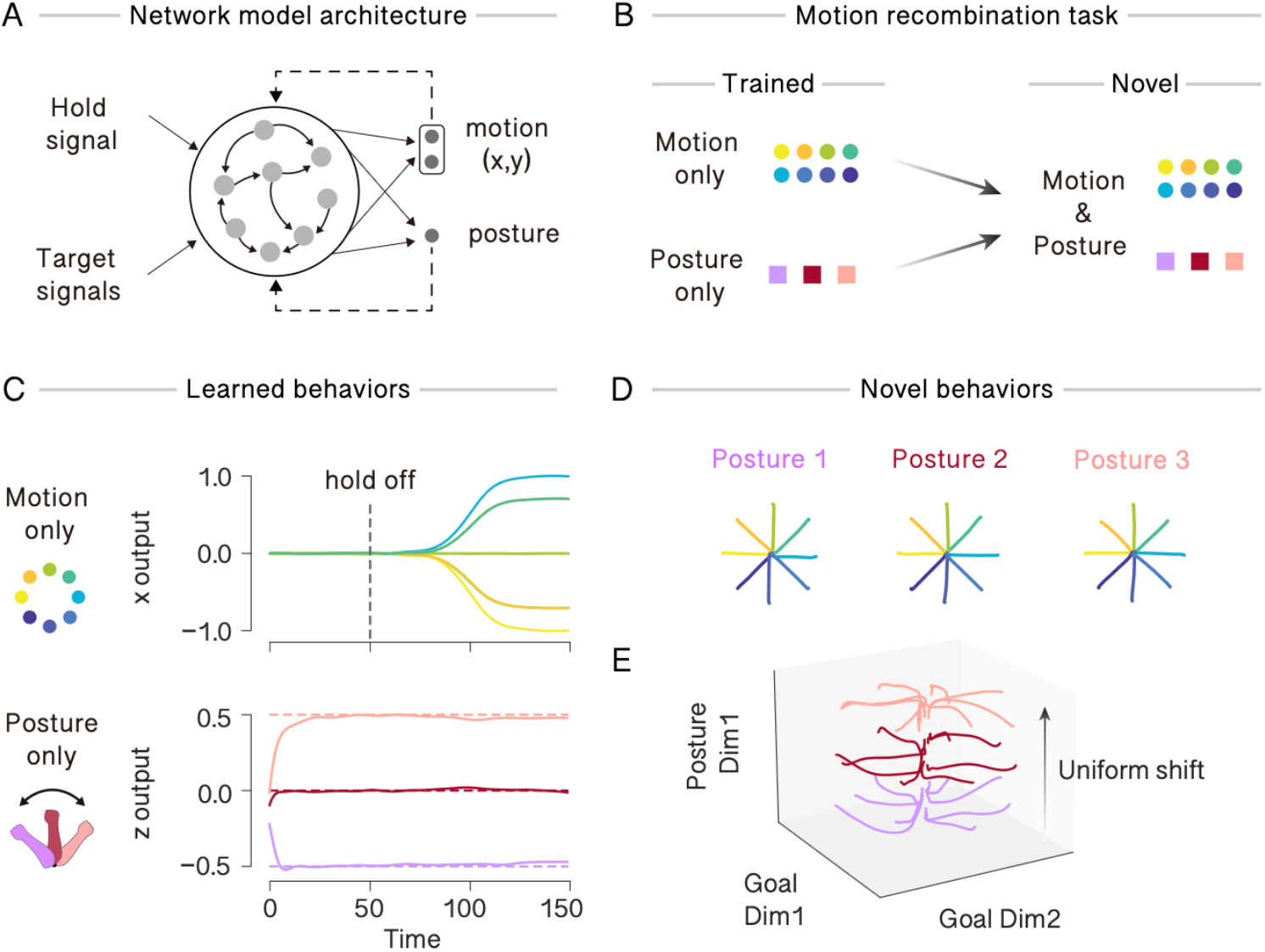
Feedback-guided learning reproduces additive reach-posture geometry. **A** Network architecture. A recurrent network receives direction-specific movement inputs and scalar posture inputs, and produces three readout channels: two movement dimensions and one posture dimension. **B** Training and test conditions. Movement and posture tasks were trained separately and were never co-presented during learning. After training, the network was tested on novel movement-posture combinations. **C** Individual task performance after separate training. Upper, x-component of the movement readout for all eight reach directions. Lower, posture readout for three posture conditions, each converging to the corresponding target level. **D** Zero-shot movement-posture composition. Combined movement readout trajectories in the two-dimensional movement plane are shown for all eight reach directions across three posture conditions. The directional structure of the reach manifold was preserved across posture conditions. **E** Combined trajectories visualized in a feedback-defined three-dimensional coordinate system. The three sets of trajectories formed approximately parallel layers offset along the posture axis, reproducing additive reach-posture geometry observed in motor cortex. In C–E, traces show the mean across 10 independent simulations.

We then tested zero-shot composition of reaching and posture by simultaneously presenting movement and posture inputs. The network generated reach trajectories that preserved both their directional selectivity and temporal structure across all posture conditions (Fig. 4D). To examine the resulting population geometry, we visualized population activity within a feedback-defined subspace corresponding to the two movement dimensions and the posture dimension. Combined reach-posture trajectories formed approximately in parallel subspaces, in which the dynamical structure of reaching was preserved within each subspace while posture produced a uniform shift along the posture axis (Fig. 4E). This geometric arrangement qualitatively recapitulates the additive population structure observed in motor cortex, where posture and movement are represented in separable subspaces and combined with minimal interference.

In summary, feedback-driven learning reproduced a motor cortex-like reach-posture geometry without requiring exposure to combined reach-posture conditions during learning. The same principle that enabled zero-shot composition in the abstract sine-wave generation task also generated an experimentally observed form of additive population geometry. These results suggest that feedback geometry provides a unifying mechanism linking the acquisition of independent computations to their later composition in novel tasks.

### Feedback-defined subspaces support zero-shot composition of higher-dimensional motor primitives

We next asked whether the same principle extends beyond low-dimensional temporal outputs to more complex, high-dimensional behaviors. To test this, we constructed a multi-output task in which a recurrent network learned three distinct coordinated dynamical primitives: an arms-up movement, a jump, and a rhythmic sway (Fig. 5A, Supplementary Movie 1). These primitives were not intended as detailed biomechanical models, but rather as test cases requiring learned computations to generate coordinated activity across multiple output dimensions. Networks were trained on each primitive separately, with each primitive associated with a distinct feedback signal, and were never exposed to combinations of primitives during training. After learning, the network generated each primitive as a coordinated sequence of multi-dimensional outputs, visualized as snapshots (Fig. 5B).

**Figure 5.**
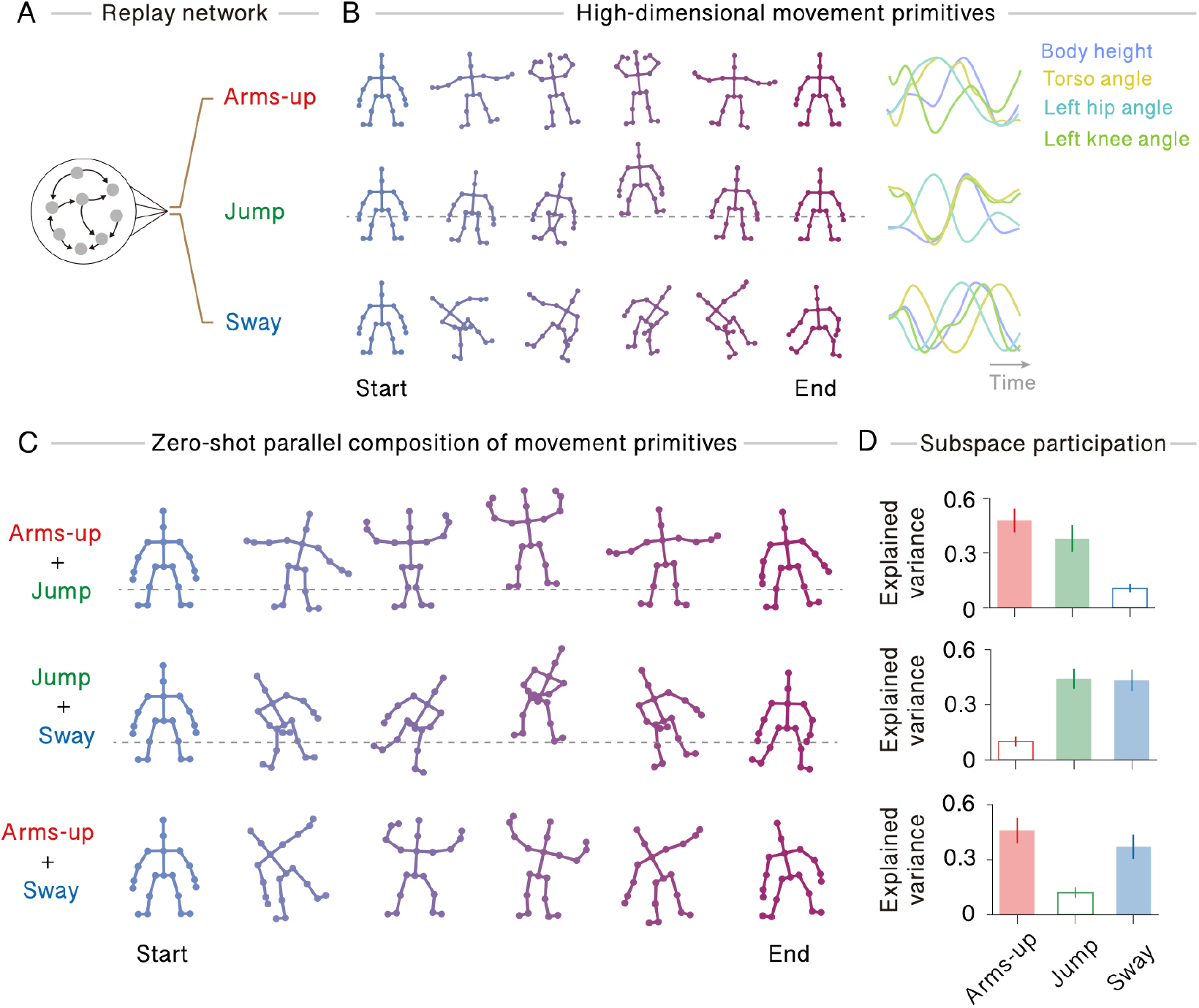
Zero-shot composition of coordinated movement-like primitives. **A** A multi-output recurrent network was trained on three movement-like primitives. Arms-up, jump, and sway were trained separately, each with a distinct feedback signal. **B** Stick-figure snapshots of the three learned primitives. Six snapshots are shown for each primitive. **C** Zero-shot composition of movement-like primitives. Pairwise combinations of primitive inputs generated coherent composite movements for arms-up and jump, jump and sway, and arms-up and sway, despite the network never being trained on these combinations. **D** Recruitment of primitive-specific subspaces during zero-shot composition. Population activity during each combined-input condition was projected onto the subspaces defined by the individual primitives. Each composition preferentially recruited the subspaces corresponding to its constituent primitives, whereas the absent primitive subspace was weakly recruited.

We then tested whether independently learned primitives could be expressed together by presenting pairs of primitive inputs simultaneously. The network generated coherent composite behaviors across all tested primitive pairs, including arms-up and jump, jump and sway, and arms-up and sway (Fig. 5C, Supplementary Movie 2). Thus, the network generated novel coordinated movements through composition of independently acquired motor primitives.

We next examined whether these composite outputs reflected selective recruitment of the corresponding primitive-specific subspaces. As in the previous analysis, we defined a subspace from the population activity evoked by each primitive presented alone and projected combined-input activity onto these subspaces (Supplementary Fig. 4). For every pairwise combination, activity was more strongly projected onto the subspaces corresponding to the presented primitives than onto the subspace corresponding to the absent primitive (Fig. 5D). Thus, zero-shot composite movements were accompanied by selective reuse of the learned primitive-specific dynamical subspaces.

In summary, feedback-driven learning enabled zero-shot composition in a multi-output setting. Novel composite outputs emerged through selective recruitment of subspaces associated with the constituent primitives, whereas subspaces corresponding to absent primitives remained weakly recruited. Therefore, the feedback-geometry principle generalized from abstract temporal patterns and reach-posture composition to the compositional reuse of coordinated dynamical primitives distributed across multiple output dimensions.

## Discussion

We show that the geometry of neural population activity reflects not only what a circuit has learned, but also how the learned computations can later be reused. Networks with matched individual task performance differed in their ability to express those tasks together, demonstrating that task acquisition and future combinability are distinct computational properties. In our model, this dissociation was controlled by learning-related feedback geometry. Distinct feedback vectors embedded task-specific dynamics into separable subspaces that supported later simultaneous expression, whereas aligned feedback embedded them in overlapping subspaces that preserved individual task performance while impairing zero-shot composition.

These findings suggest a broader functional role for neural manifolds. Separable (or orthogonal) subspaces are often viewed as a mechanism for representing multiple variables within the same population while minimizing interference^14,32–34^. Our results suggest that such geometry can preserve computations as reusable dynamical components for future behavior. In successful networks, combined inputs did not trigger switching between previously learned trajectories. Instead, they generated a single population trajectory whose projections onto task-specific subspace recovered the independently learned dynamics. Thus, manifold geometry served not merely as a representational scaffold but as a substrate for compositional computation.

The comparison with BPTT-trained networks further supports this interpretation. Standard RNNs trained on the same individual tasks learned the component mappings accurately but generalized poorly to the combined condition^35^. Thus, successful zero-shot composition cannot be explained by task acquisition alone. Instead, it depends on how learning embeds computations within the recurrent state space. The systematic feedback-angle manipulation strengthened this conclusion (Fig.3D). Performance remained high when feedback vectors were sufficiently separated but declined as they became aligned, indicating that feedback geometry continuously controls the degree to which learned dynamics can be expressed in parallel.

The population-level mechanism identified here may map directly to experimentally observed neural geometry in motor cortex. During reaching from different postures, movement-related dynamics and posture-related activity occupy approximately orthogonal subspaces, allowing posture to shift the reach manifold without distorting its internal structure^15^. Our model reproduced this additive reach-posture organization despite never being trained on combined reach-and-posture conditions. This suggests that such geometry may reflect not only the statistics of experienced movements but also the structure of plasticity signals that shape circuit dynamics during learning. The same principle extends to coordinated movement-like primitives, where novel composite outputs emerge through selective reactivation of primitive-specific dynamical subspaces^36^.

These results make several experimental predictions. At the population level, tasks acquired under distinct feedback, teaching, reward, or contextual signals should become embedded in more separable subspaces than tasks acquired under shared feedback signals^37^. This separation should predict future combinability more strongly than single-task performance. At the circuit level, disrupting a task-specific feedback pathway during learning should impair later zero-shot composition without necessarily affecting performance of the individual task. More generally, the model predicts that the geometry of learning-related feedback pathways constrains which learned computations can later be jointly expressed.

Several limitations remain. The tasks studied here involve additive forms of parallel composition, and the networks rely on linear readouts with externally specified input combinations. More complex forms of compositional behavior, including conditional routing, multiplicative interactions, or hierarchical sequencing, may require additional mechanisms beyond the principle demonstrated here. In addition, the feedback vectors in our model were preconfigured rather than learned. Biophysically, such vectors may correspond to projection patterns from upstream areas^38–41^. Thus, our model isolates the computational consequence of feedback geometry, rather than its biological origin. How such feedback pathways are acquired or reorganized to support future combinability remains an open question.

Together, our results suggest that learning-related feedback signals shape recurrent dynamics in ways that determine whether independently acquired computations can later be expressed together.

## Methods

### The neural network model

We used recurrent networks of rate units. The membrane potential of unit *i* at time *t* was denoted by *x*_*i*_(*t*), and the firing rate was defined as *r*_*i*_(*t*) = tanh*x*_*i*_(*t*). In vector form, the network dynamics were described by

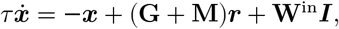

where ***x*** is the membrane-potential vector, ***r*** = tanh(***x***) is the firing-rate vector, **G** is a fixed random recurrent matrix, **M** is a plastic recurrent matrix, **W**^in^ is the input weight matrix, and ***I*** is the external input vector. Unless otherwise stated, *τ* = 10 ms.

The fixed recurrent matrix **G** was initialized from a zero-mean Gaussian distribution with variance scaled by network size. In densely connected simulations, entries of **G** were drawn with standard deviation 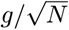. In the sparse whole-body movement simulation, nonzero entries were drawn with standard deviation 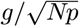, where *p* is the connection probability. The plastic recurrent matrix **M** was initialized to zero in the temporal-generation and reach-posture simulations, and to small random values in the whole-body movement simulation. The initial network state was sampled once from a Gaussian distribution and reused across trials within each simulation.

We used linear readouts as network output units. For a scalar-output task, the readout was

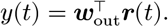

For multi-output tasks, each output dimension *a* had an independent linear readout,

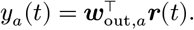

### The learning rule

Recurrent synapses were trained using a predictive alignment plasticity rule (Asabuki and Clopath, 2025). The rule updates the plastic recurrent matrix **M** so that the plastic recurrent drive **M*r***(*t*) matches a feedback-defined reference drive. For a scalar output, the reference drive was

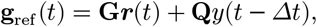

where **Q** is a fixed feedback vector, of which the elements were independently sampled from a uniform distribution with range set by Q_scale_. The recurrent update was

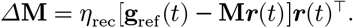

For multi-output tasks, the feedback term was the sum of output-specific feedback vectors weighted by the previous readout activity,

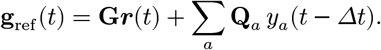

Readout weights were trained by a delta rule,

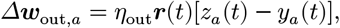

where *z*_*a*_(*t*) is the target output. Unless otherwise stated, *η*_rec_ = *η*_out_ = 10^−3^. In the whole-body movement simulation, both learning rates were 0.3 × 10^−3^ . During testing, all recurrent and readout weights were fixed.

### Temporal-generation task

For the minimal zero-shot parallel-composition task in Figs. 2 and 3, the network received two constant input signals, *S*_1_ and *S*_2_. Each input was associated with a different sinusoidal target output. The first target completed one cycle over the trial and the second target completed two cycles over the trial,

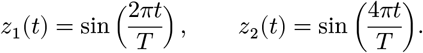

Each target was z-scored before training. The two tasks were trained separately. On alternating training episodes, the network received either *S*_1_ alone and was trained to output *z*_1_(*t*), or *S*_2_ alone and was trained to output *z*_2_(*t*). The network was never trained with *S*_1_ and *S*_2_ presented together.

We compared two feedback conditions. In the different-feedback condition, the two tasks were trained with independent feedback vectors **Q**_1_ and **Q**_2_ . In the same-feedback condition, the second task used a copy of the first feedback vector. After training, zero-shot composition was tested by presenting both inputs simultaneously. The target for this untrained condition was defined as

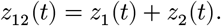

No recurrent or readout weights were updated during this test phase. Performance was quantified as the Pearson correlation between the z-scored network output and the z-scored combined target. In the simulations used for Figs. 2 and 3, *N* = 600, *T* = 100, *g* = 1.5, Q_scale_ = 2.0, and networks were trained for 1,000 episodes. Statistics were computed across 20 random seeds.

### Manifold and subspace analysis

To analyze population geometry in Figure 3, recurrent activity was recorded during the two single-input conditions and the combined-input condition. For each task, a task-specific activity subspace was obtained by applying principal component analysis to the corresponding single-task activity matrix after subtracting the time-averaged activity vector. Unless otherwise stated, the first 10 principal components defined each task subspace.

Let *U*_1_ and *U*_2_ denote the orthonormal bases of the subspaces obtained from the *S*_1_-alone and *S*_2_-alone trajectories. Before projection, each trajectory was centered by subtracting its own time-averaged population activity vector. For example, let 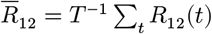 denote the mean activity vector of the combined-input trajectory. The centered combined trajectory was then projected onto each task subspace as

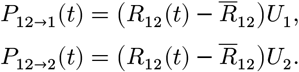

Projection similarity was quantified by the Pearson correlation between matched principal-component time courses. For each network, correlations were averaged over the first three principal components.

Subspace orthogonality was quantified by principal angles between *U*_1_ and *U*_2_. If *σ*_*k*_ are the singular values of *U*^⊤^*U*, the principal angles were defined as

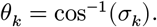

The mean principal angle was used as a scalar measure of task-subspace separation. We also quantified cross-talk in the combined condition as the absolute correlation between the first projected component in Subspace 1 and the first projected component in Subspace 2.

### Reach-posture task

In Figure 4, the network produced three readout channels: two motion outputs, *x*(*t*) and *y*(*t*), and one posture output, *z*(*t*). The motion task consisted of reaches in eight directions. For reach direction *θ*, the target trajectory was

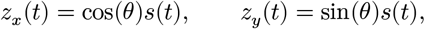

where *s*(*t*) was a sigmoid reach profile beginning after a hold period. Trials lasted *T* = 150 time steps with a hold period of 50 time steps. During the hold period, the input consisted of a hold signal plus a direction-specific motion input. After the hold period, the hold signal was removed and the direction-specific motion input remained.

The posture task required the network to maintain one of three scalar posture outputs, z∈{- 0.5, 0, +0.5}. Motion and posture trials were trained separately. During motion trials, the x and y readouts were trained to generate the reach trajectory, while the posture readout was trained toward zero. During posture trials, the posture readout was trained toward the corresponding scalar posture target, while the motion readouts were trained toward zero. Motion and posture inputs were never presented together during training. After training, we tested all combinations of eight reach directions and three posture conditions without further learning.

After training, zero-shot reach-posture composition was tested by presenting a motion input and a posture input simultaneously. No weights were updated during testing. We tested all combinations of eight reach directions and three posture conditions. Preservation of reach geometry was quantified by the correlation between the two-dimensional motion trajectory in the motion-alone condition and the corresponding two-dimensional motion trajectory in the combined condition. Population trajectories were visualized in the three-dimensional readout space defined by the *x, y*, and posture outputs.

To visualize recurrent population activity in a feedback-defined coordinate system, we projected the activity *R*(*t*) onto the normalized feedback vectors *Q*_*x*_, *Q*_*y*_, and *Q*_*z*_ . Coordinates were computed separately for each seed and then averaged across seeds. Unless otherwise stated, the raw feedback vectors were normalized but not orthonormalized, preserving their interpretation as output-specific feedback axes. The reach-posture simulations used N=600, T=150, T_hold_ =50, g=1.2, Q_scale_ =3.0, posture input scale =0.5, and 2,000 training episodes, averaged over 10 random seeds.

### Coordinated movement-like dynamics

In Figure 5, the network produced a 15-dimensional kinematic output vector consisting of root translation and parent-relative joint-angle variables. The kinematic outputs were converted into two-dimensional stick-figure poses using a forward-kinematics model. The root variables specified global hip translation, the torso variable specified the torso angle, and all other joint variables specified parent-relative angular deviations from a neutral pose. The network learned three movement primitives: arms-up, jump, and sway. Each primitive was defined as a smooth trajectory in the 15-dimensional kinematic space. The arms-up primitive included coordinated bilateral arm elevation, the jump primitive included crouch, launch, aerial, and landing phases with arms kept down, and the rhythmic sway primitive included oscillatory torso, limb, and root motion.

The whole-body network used *N* = 5,000 recurrent units, three primitive-specific inputs, and 15 output dimensions. The fixed recurrent matrix was sparse, with connection probability *p* = 0.1 and gain *g* = 1.2. Primitive input weights were drawn uniformly from [−0.1, 0.1]. Primitive-specific feedback matrices *Q*_*k*_ were drawn independently for each primitive and output dimension. The recurrent and readout learning rates were both 0.3 × 10^−3^. Each training episode contained only one primitive, and primitives were cycled across episodes. The network was trained for 1,000 episodes. After training, we tested single primitives and novel combinations without further learning. For a combination of primitives, the input was the sum of the corresponding primitive inputs. Stick-figure visualizations were generated by applying the forward-kinematics model to the 15-dimensional network output at each time step.

### Principal component analysis of the movement-like dynamics

Principal component analysis was used to define primitive-specific activity subspaces from single-primitive trials. For primitive *i*, the recurrent firing-rate trajectory was centered by subtracting its time-averaged activity, and the leading principal components defined the subspace *U*_*i*_, of which the columns were orthonormal. Single-primitive and combined-input trajectories were then projected onto these subspaces to assess whether combined inputs reactivated the dynamics associated with each component primitive.

Because primitive subspaces could contain both primitive-specific and shared temporal structure, we estimated and removed a shared component using only single-primitive activity. For each primitive subspace, we constructed the projection matrix

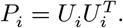

We then computed the mean projection matrix across the *K* primitives,.

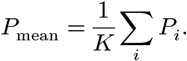

Shared directions were identified by eigendecomposition of the mean projection matrix,

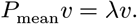

For a unit-norm eigenvector *v*, the eigenvalue is

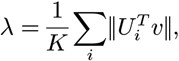

and therefore measures how strongly the direction *v* is represented across primitive subspaces. A shared subspace *U*_sh_ was constructed from eigenvectors with *λ* > 0.8 for the analysis in Fig. 5 and removed from each primitive subspace by orthogonal projection,

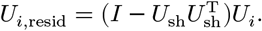

The columns of *U*_*i*,resid_ were then orthonormalized, and the resulting residual subspaces were used for the trajectory projections and variance-explained analysis in Fig. 5. This procedure did not use information about which primitives were present in the combined conditions.

### Simulation details

All simulations were performed in custom Python code using NumPy, SciPy, scikit-learn, Numba, and Matplotlib. Differential equations were numerically integrated with an Euler method with an integration time step of 1 ms. Random seeds were sampled independently for each network replicate.

## Data availability

All numerical datasets necessary to replicate the results shown in this article can easily be generated by numerical simulations with the software code available upon publication. No datasets were generated during this study.

## Code availability

Example source codes used for the present numerical simulations and data analysis will be made available upon publication.

## Author Contributions

Y.O. and T.A. conceived the study. T.A. designed the model, developed the mathematical analysis, and supervised the project. Y.O., A.A., and T.A. performed the simulations and numerical analyses. All authors interpreted the results and wrote the manuscript.

## Acknowledgments

This work was supported by RIKEN Center for Brain Science (TA) and RIKEN Cluster for Pioneering Research (TA). T.A. thanks Zihan Liu for fruitful discussions and helpful comments on the manuscript.

## Supplementary Figures

**Supplementary Figure 1.**
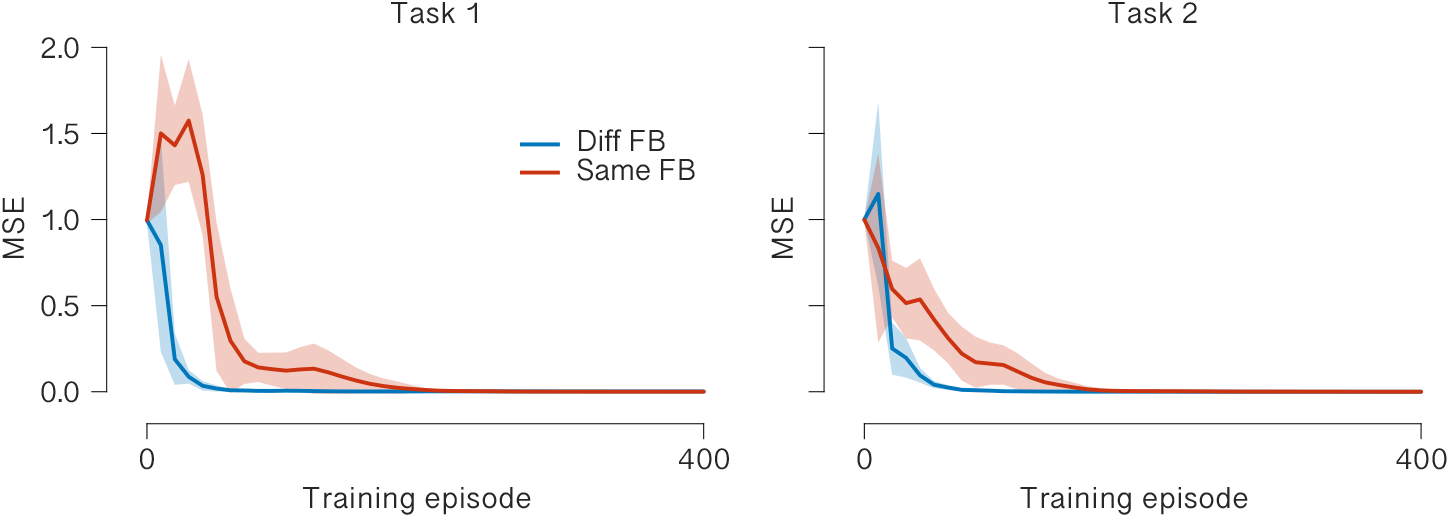
Learning curves for the two individual tasks. Learning curves for Task 1 and Task 2 under the different-feedback and same-feedback conditions. Lines show the mean across 20 independent simulations, and shaded regions show s.d. Both conditions eventually acquired each individual task with low error.

**Supplementary Figure 2.**
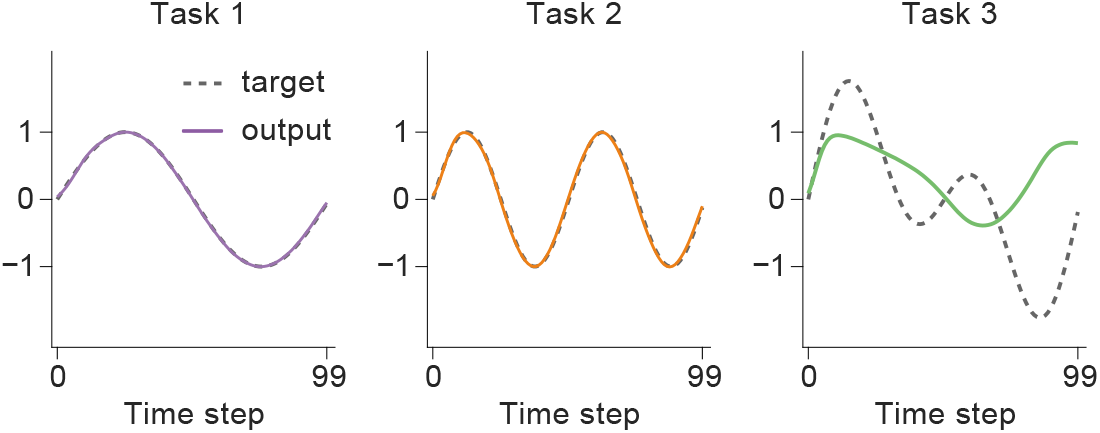
BPTT-trained networks learn individual tasks but fail to recombine them. Example output traces from a standard recurrent network trained by backpropagation through time on the two individual temporal-generation tasks. The network accurately reproduced the target output during Task 1 and Task 2 when each input was presented alone. However, when the two inputs were presented simultaneously, the output failed to match the untrained combined target.

**Supplementary Figure 3.**
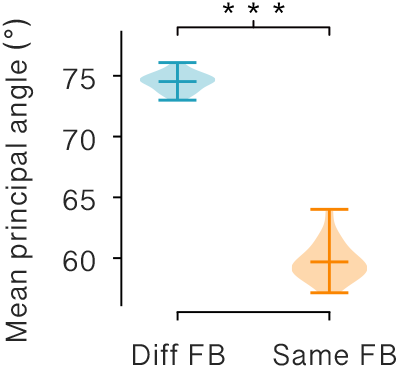
Different feedback vectors produce more separated task subspaces. Mean principal angles between task-specific activity subspaces in the different-feedback and same-feedback conditions. Task subspaces were defined from recurrent activity during the two single-task conditions.

**Supplementary Figure 4.**
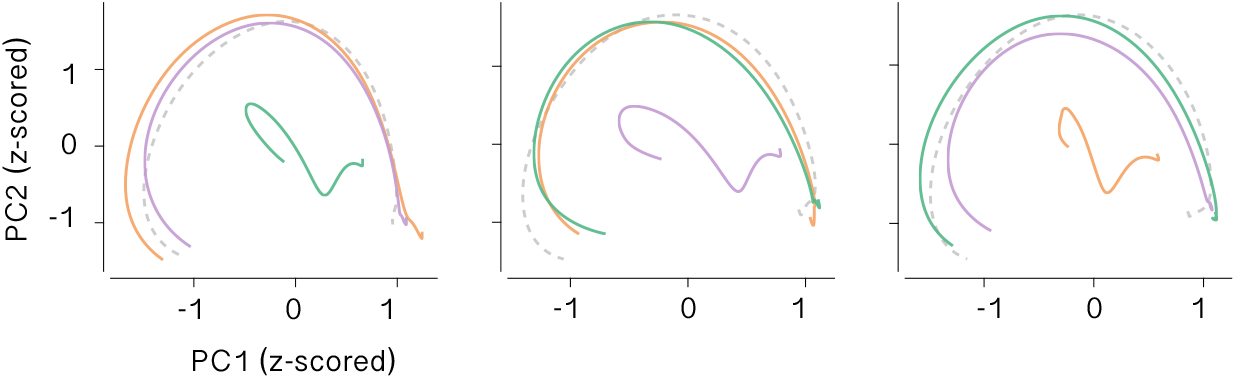
Combined movement-like inputs selectively reactivate primitive-specific trajectories. Principal component projections of recurrent activity for the higher-dimensional movement-like task. Each panel shows activity projected onto one primitive-specific subspace, defined from the corresponding single-primitive trajectory.

## Supplementary Information

### Low-rank mean-field theory of zero-shot parallel composition

We developed a low-rank mean-field description of the trained recurrent network^1^. Our prior work has shown that each learned computation is represented by a structured low-rank mode embedded in the recurrent connectivity^2^.

Our analysis relies on a Dynamical Mean-Field Theory (DMFT). Intuitively, distinct feedback vectors enhance the two learned computations to be embedded in separate directions of the recurrent dynamics. As a result, the network can activate one computation without substantially disrupting the other. In mean-field terms, this corresponds to statistically orthogonal low-rank modes whose collective variables are approximately decoupled, allowing the zero-shot composition to drive both computations in parallel. By contrast, aligned feedback vectors embed the two computations in overlapping modes, producing off-diagonal coupling between collective variables and thereby causing cross-talk during combined-input conditions.

We consider a rate recurrent network of *N* units,

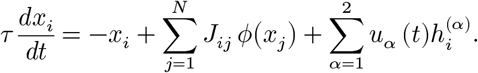

Here, *x*_*i*_ is the membrane potential of unit *i, ϕ*(*x*_*i*_) is its firing rate, *u*_*α*_(*t*) is the input associated with task *α*, and *h*^(*α*)^ is the corresponding input vector. We approximate the trained recurrent connectivity as the sum of a random recurrent component and a structured low-rank component^1^,

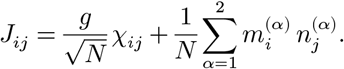

Here, *χ*_*ij*_ are independent random variables with zero mean and unit variance, *g* controls the strength of the random recurrent connectivity, and *m*^(*α*)^, *n*^(*α*)^ are the right and left connectivity vectors of the learned mode associated with task *α*. In this description, task 1 and task 2 are embedded as two structured connectivity modes within the same recurrent population.

We define the collective variable associated with mode *α* as

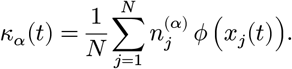

Under this definition, the structured part of the recurrent input can be rewritten as

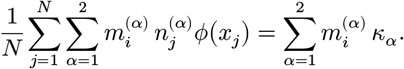

Therefore, the network dynamics can be expressed as

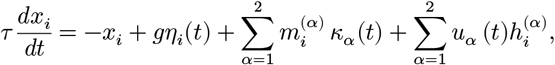

where

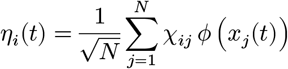

is the effective input generated by the random recurrent component^3^. In the large *N* limit, DMFT theory treats *η*_*i*_(*t*) as a Gaussian process whose temporal covariance is determined self-consistently by the population activity,

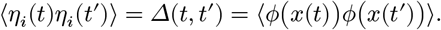

The random component therefore contributes a high-dimensional fluctuating background, whereas the low-rank component generates low-dimensional collective dynamics through *κ*_1_ and *κ*_2_.

We next derive the effective coupling between the two collective variables. Starting from the definition of *κ*_β_,

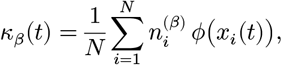

and substituting the mean-field expression for *x*_*i*_(*t*), we obtain the self-consistent relation

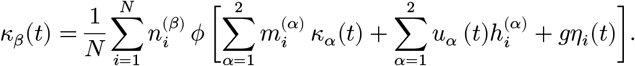

In the large *N* limit, this empirical average can be replaced by an average over the joint distribution of the loadings *n*^(*β*)^, *m*^(*α*)^, *h*^(*α*)^, and the Gaussian field *η*,

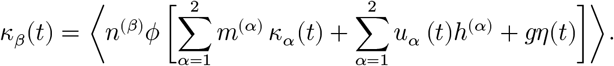

Equivalently, the collective dynamics can be written as

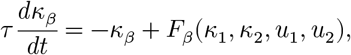

where

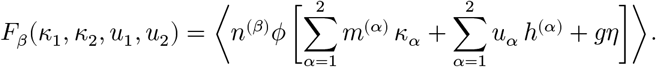

To identify the terms that control interference between the two learned computations, we linearize these collective dynamics around a background trajectory 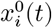. Let

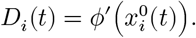

For small perturbations in *κ*_*α*_ and *u*_*α*_, we use

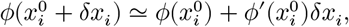

with

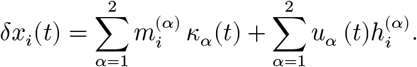

Substituting this expansion into the definition of *κ*_*β*_ gives

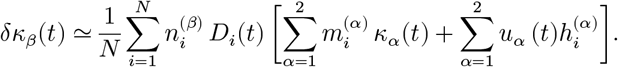

Therefore,

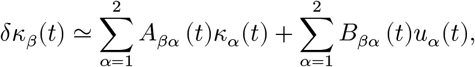

where the effective recurrent coupling matrix is

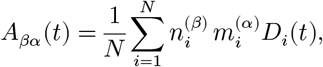

and the effective input coupling matrix is

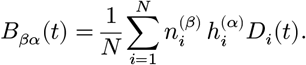

Thus, the linearized collective dynamics are

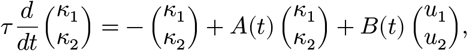

with

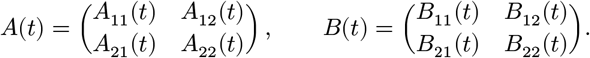

This expression makes the condition for zero-shot parallel composition explicit. The diagonal terms *A*_11_ and *A*_22_ describe the self-dynamics of the two learned modes. In contrast, the off-diagonal terms *A*_12_ and *A*_21_ describe cross-coupling between the two computations. For example, *A*_12_ measures the extent to which activation of mode 2 changes the dynamics of mode 1. Similarly, *B*_12_ and *B*_21_ measure cross-drive of one collective variable by the input associated with the other task.

The key question is therefore when the off-diagonal terms are negligible. For *α* ≠ *β*, define

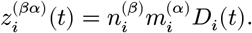

Then

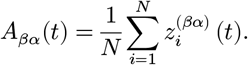

If the learned modes are statistically orthogonal, centered, and not systematically correlated through the gain factor *D*_*i*_(*t*), then

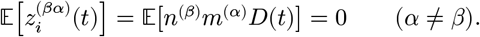

Under these conditions,

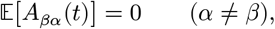

and

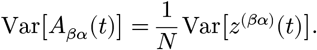

Therefore, in the large-*N* limit,

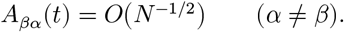

Thus, the off-diagonal recurrent couplings vanish up to finite-size fluctuations when the learned low-rank modes are statistically orthogonal. An analogous argument applies to the off-diagonal elements of the input coupling matrix. In our model, the input vectors are fixed and are not learned. Thus, we do not assume that the input weights are explicitly aligned with the learned modes. Instead, the relevant condition is that each fixed task input has an effective projection onto the mode formed during the corresponding task, while its projection onto the other task mode remains small. Because the two input vectors are independently drawn and the two learned modes are shaped under distinct feedback conditions, the cross-input couplings are expected to vanish in the large *N* limit,

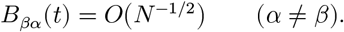

The diagonal terms *B*_*αα*_, in contrast, need not vanish, because the learned mode *n*^(*α*)^ is formed from activity evoked under the corresponding input condition *h*^(*α*)^.

In this regime, the effective collective dynamics are approximately block diagonal,

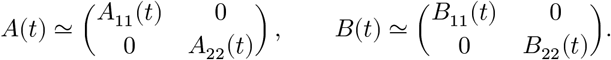

The two collective variables therefore obey approximately decoupled dynamics,

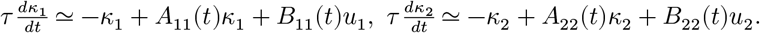

Consequently, input *u*_1_ primarily drives *κ*_1_, and input *u*_2_ primarily drives *κ*_2_. When both inputs are presented simultaneously, the two collective variables are co-activated but remain weakly coupled,

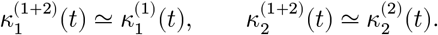

If the output is read from the same low-dimensional structure,

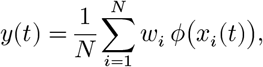

and the readout has components aligned with the two learned modes, then to leading order,

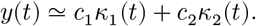

Therefore, under simultaneous input,

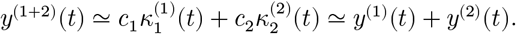

This gives a mean-field explanation of zero-shot parallel composition. Distinct learned modes define approximately independent collective variables. Simultaneous inputs co-drive these variables, and the output combines their contributions additively.

The same analysis explains why aligned feedback vectors impair composition. When feedback vectors are aligned, learning is expected to embed the two tasks in overlapping modes rather than statistically orthogonal modes. In that case,

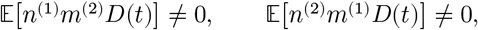

and therefore

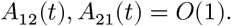

The effective dynamics are no longer block diagonal,

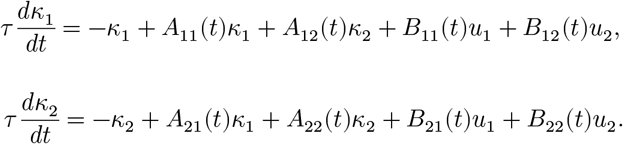

Now, activation of one learned component changes the dynamics of the other through the cross-coupling terms *A*_12_*κ*_2_ and *A*_21_*κ*_1_. The combined-input response is therefore no longer a simple superposition of the two single-input responses. This provides a mechanism of why aligned feedback can support individual task learning while impairing zero-shot composition.

This analysis also explains the feedback-angle dependence observed in Fig. 3D. When feedback vectors are partially aligned, learning is expected to embed the two tasks in modes with nonzero statistical overlap, causing off-diagonal couplings to scale with the degree of feedback alignment rather than vanishing. Importantly, composition is not expected to fail as soon as the feedback vectors deviate from perfect orthogonality. As long as the induced off-diagonal couplings remain small relative to the diagonal self-couplings and the stability margin of each learned mode, their effect remains perturbative, and simultaneous inputs can still drive the two collective variables approximately independently. However, when the feedback angle becomes sufficiently small, the induced cross-coupling can become comparable to this mode-separation margin, causing the combined-input response to deviate rapidly from the sum of the two single-input responses. This provides a mean-field interpretation of the crossover observed in Fig. 3D, where composition remains relatively robust over intermediate angles but deteriorates rapidly once feedback vectors become too aligned.

In summary, zero-shot parallel composition can be understood as a consequence of the effective block diagonalization of low-dimensional collective dynamics. Distinct feedback vectors promote statistically orthogonal learned modes, causing the off-diagonal couplings between collective variables to vanish as *O*(*N*^−1/2^). Aligned feedback vectors generate overlapping modes, producing finite off-diagonal coupling and interference.

